# Strategic template filtering accelerates fragment-based peptide docking

**DOI:** 10.64898/2026.03.26.714397

**Authors:** Nirit Trabelsi-Mescheloff, Julia K. Varga, Alisa Khramushin, Sergey Lyskov, Ora Schueler-Furman

**Author notes:** These authors contributed equally (Nirit Trabelsi-Mescheloff, Julia K. Varga).

## Abstract

Peptide-protein interactions are often transient and structurally elusive, necessitating computational approaches to identify both binding sites and peptide conformations. PatchMAN, one of the leading but computationally expensive biophysic-based global peptide-docking protocols, addresses this challenge by treating peptide docking as a protein-folding problem, using structural motifs from solved *structures* as templates that are subsequently refined using Rosetta FlexPepDock. Here we present PatchMAN2, which introduces 1) strategic fragment filtering and 2) local docking modes that focus sampling on relevant surfaces or known binding regions, thereby reducing the high computational cost of the original implementation due to extensive refinement of many non-productive low-quality fragments. Benchmarking shows that PatchMAN2 removes ∼30-70% of unnecessary fragments while preserving accuracy, substantially reducing runtime and improving the practical efficiency of peptide-protein docking.

## 1. Introduction

Peptide-protein interactions are often transient and weaker than domain-domain interactions involving well-defined folded structural units, making them well suited for rapid regulation of cellular processes. It is estimated that cells harbor approximately one million peptide-protein interactions^1^, yet experimentally solved structures exist for only a small fraction of them. As a result, computational modeling is essential for studying their atomic details and elucidating key contributions to binding. A central challenge arises from peptide flexibility that needs to be taken into account together with orientation of the peptide on the receptor surface: the peptide conformation prior to association is typically unknown, requiring identification of both the binding site and the bound peptide conformation at the same time. Despite the rapid expansion of deep-learning-based docking protocols^2–7^, these approaches still do not always succeed, particularly for interaction types that are underrepresented in training datasets like peptide-protein interactions^8–11^. In fact, the search space can be reduced since the binding site can often be reliably determined from experimental evidence such as mutational scanning, or inferred computationally based on homologous interaction structures, solvent mapping^12,13^, conserved interface residues^14^ or other methods^15–19^. Incorporating this information can substantially reduce the docking search space and the associated computational cost.

Several workflows have been developed along the years both for global and local docking, often relying on explicit biophysical knowledge to constrain sampling^20–25^. One effective strategy is based on the observation that peptide-protein interactions can be viewed as a special case of protein monomer folding, in which the peptide complements the globular fold of the receptor and reuses structural motifs already present in monomer structures^26^. Based on this concept, our group developed the global peptide docking protocol PatchMAN^27^. PatchMAN separates the receptor surface into patches and searches for similar structural motifs in solved structures, from which complementary fragments are extracted and threaded with the peptide sequence to generate peptide-receptor templates. These are then refined using Rosetta FlexPepDock (FPD)^28^. The top 1% scoring models are clustered, and the top 10 best-scoring cluster representatives returned as final predictions. In benchmarking against other methods, PatchMAN ranks among the most accurate currently available global docking protocols^27^, with similar performance relative to AlphaFold2 (AF2)^29^. Because PatchMAN does not rely on sequence similarity during the search, it can extract templates from structures with low sequence identity or even unrelated folds, enabling straightforward extension to peptide design.

Although PatchMAN demonstrates promising performance, its computational cost is dominated by full-atom refinement of a large number of fragments using FPD. Removing unproductive fragments prior to refinement could therefore substantially accelerate and improve the protocol. Here we present PatchMAN2, an enhanced version that introduces filtering strategies to focus sampling on promising candidates. In particular, a buried surface area (BSA)-based filter eliminates fragments unlikely to yield native-like conformations. In addition, non-binding surfaces (such as obligatory multimer interfaces) can be masked. Finally, when binding sites are known or can be predicted, sampling can be restricted to relevant regions. By concentrating computational effort on the relevant search space, PatchMAN2 substantially reduces runtime and resource usage, while allowing improved accuracy relative to the original protocol. PatchMAN2 is available on GitHub (https://github.com/Furman-Lab/PatchMAN) and as a web server on the ROSIE platform^30^ (https://r2.graylab.jhu.edu/apps/submit/patchman).

## 2. Materials and Methods

### 2.1. Data Sets

*Benchmarks:* We used two data sets of peptide-protein complex structures from our previous studies: (1) The PFPD (PiperFlexPepDock) set^25^ contains 24 complexes (we removed from the original PFPD set of 26 complexes two Protein Data Bank, PDB^31^, IDs: for 2FMF^32^, the peptide structure is possibly heavily influenced by crystal contacts; for 1NVR^33^, the peptide side chains are unresolved according to the study that reports this structure); (2) the LNR (Large Non Redundant) set^29^ contains 96 complexes. Both datasets are non-redundant at the receptor ECOD^34^ domain level to provide unbiased assessment of docking protocols. For the LNR dataset, we used only a subset of proteins for which unbound structures are available in the PDB (as in the original PatchMAN implementation^27^). These were used as input for all PatchMAN runs (see **Supplementary Table S1**).

*Homolog complex selection:* For each entry in the datasets we selected the structure of a random homolog complex. Homolog complexes were defined based on the ECOD classification of the receptor domain, with a peptide located within 6 Å RMSD of the peptide in the benchmark structure (see **Supplementary Table S1**).

### 2.2. Measure of Performance

*Interface residues* were defined as residues whose C_β_ atoms are within 8 Å of a C_β_ atom in another chain (r_Cβ-Cβ_ ≤ 8 Å; C_α_ for glycine). Root mean square deviation, *RMSD,* was calculated over the peptide interface residue backbone atoms (N, C_ɑ_, C, O), comparing the native structure to the models.

*Successful models* were defined as complexes modeled within 2.5 Å RMSD from the native structure. *PatchMAN docking performance* was assessed based on the best RMSD among the top 10 cluster representatives after clustering, as in previous studies^25,27^.

### 2.3. Filters

#### 2.3.1 BSA filtering

The buried surface area (BSA) was calculated as:

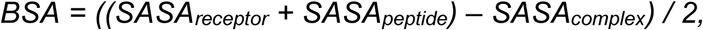

where solvent accessible surface area (SASA) values were computed separately for the receptor, peptide, and complex. Of note, while our initial analysis was performed using freeSASA^35^ to calculate the buried surface, in PatchMAN2 we implemented BSA calculation using PyRosetta^36^. Both methods are in good agreement, with a small shift in the calculated values (see **Supplementary Table S2**).

*Definition of BSA cutoff:* Because the distribution of peptide lengths is uneven we concatenated 4 consecutive lengths (e.g. 3-6 amino acids, 7-10 amino acids, *etc.*) for cutoff calculation. To ensure that we prevent filtering of near-native conformations, we subtracted 50 Å^2^ from the minimal value. The final rule for cutoff derivation is:

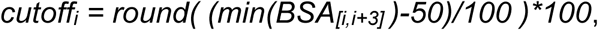

where *i* is the peptide length reflecting the cutoff for the 4-residue length interval [*i, i+3*].

#### 2.3.2 Definition of area of interest

*Definition of mask area*: For complexes forming obligatory homomultimer interfaces, the mask was defined by selecting receptor residues at the interface with another chain in the native complex structure (r_Cβ-Cβ_ ≤ 8 Å).

*Definition of focus area based on homolog structures:* The focus area on the receptor was defined as residues within 6 Å heavy atom distance from the peptide in the homolog complex after superposition of the homolog domains.

*Definition of hotspot residues:* We used a local installation of the Robetta alanine scanning protoco^l37^ to identify hotspots in the receptor of the native crystal structures. The top 3 residues with the strongest impact upon mutation were selected as hotspots (see **Supplementary Table S1**). To define the *hotspot area*, we extend the area around these hotspots to include receptor residues within 8 Å ≤ r_Cβ-Cβ_ (C_α_ for glycine), where r_Cα-Cα_> r_Cβ-Cβ_, to ensure that residues point in the direction of the binding site.

### 2.4. Run command lines

The following commands were used to run the different pipelines (important specific flags are highlighted):

BSA filtering (default run in the updated protocol):

~~~
python3 PatchMAN_protocol.py -a <NATIVE_PDB> -m -b
<BENCHMARK_FILE> <UNBOUND_PDB> <PEPTIDE_SEQUENCE>
~~~

Mask (default mode invoked with either of the -s or -l flags):

~~~
python3 PatchMAN_protocol.py -a <NATIVE_PDB> -m **-s** <MASK_PDB> -b
<BENCHMARK_FILE> <UNBOUND_PDB> <PEPTIDE_SEQUENCE>
~~~

Focus:

~~~
python3 PatchMAN_protocol.py -a <NATIVE pdb> -m **-f** -s <FOCUS_PDB>
-b <BENCHMARK_FILE> **-t 3** <UNBOUND_PDB> <PEPTIDE_SEQUENCE>
~~~

Hotspot:

~~~
python3 PatchMAN_protocol.py -a <NATIVE_PDB> -m **-o** -s
<HOTSPOTS_PDB> -b <BENCHMARK_FILE> **-t 3** <UNBOUND_PDB>
<PEPTIDE_SEQUENCE>
~~~

Supplying a list of residues either with *-s* or *-l* invokes the filter mode: *-s <SPECIAL_PDB>* provides a pdb file containing the residues of the mask/focus/hotspot region (this file must use the same residue numbering as the *<UNBOUND_PDB>*), while *-l* can be used to include a list of residues. The default is to mask the provided residues. This can be changed by adding *-f* for the focus mode or *-o* for the hotspot mode.

Further input parameters: *-a <NATIVE_PDB>* refers to the solved structure (in a benchmark run. The RMSD to this structure will be calculated to assess performance); *-m* refers to receptor backbone minimization during FPD refinement; *-b <BENCHMARK_FILE>* refers to a list of complexes with the same receptor as the input (these excluded from the MASTER run, to provide a real-world scenario in which the structure has not been solved yet; see **Supplementary File S1**); *<UNBOUND_PDB>* indicates the input structure of the receptor, and *<PEPTIDE_SEQUENCE>* the input sequence of the peptide. *-t* indicates the number of models to generate with the FlexPepDock refinement protocol for each fragment.

### 2.5. Software

The modifications were implemented in the PatchMAN pipeline, using python3.11, and PyRosetta release 2025.3^36^. The updated protocol PatchMAN2 is available on Github (https://github.com/Furman-Lab/PatchMAN/releases/tag/v2.2). For easier installation, refinement and clustering by command line, Rosetta commands from the original workflow were entirely replaced with the same refinement and clustering protocols implemented in PyRosetta. Analysis and plots were performed using python3.11, and protein structures were visualized using ChimeraX v1.8^38^.

## 3. Results

The PatchMAN docking pipeline consists of three main stages. First, the receptor surface is divided into structural patches that serve as queries for identifying similar motifs in databases of solved protein structures. Second, peptide template fragments are extracted from structural matches identified by the MASTER^39^ search and threaded with the peptide sequence. Finally, the resulting templates are refined using Rosetta FlexPepDock (FPD) to generate high-resolution peptide-protein models. PatchMAN2 introduces several filtering strategies that remove unpromising candidates during the first two stages of the pipeline, thereby reducing the number of fragments entering the computationally expensive refinement stage. Specifically, filters can be applied during **patch selection** (masking irrelevant regions or focusing on known binding areas; **Fig. 2** and **Fig. 3**) and/or during **template extraction** (e.g., filtering fragments based on buried surface area; **Fig. 1**). The following sections describe these strategies and evaluate their impact on docking performance and runtime.

**Figure 1.**
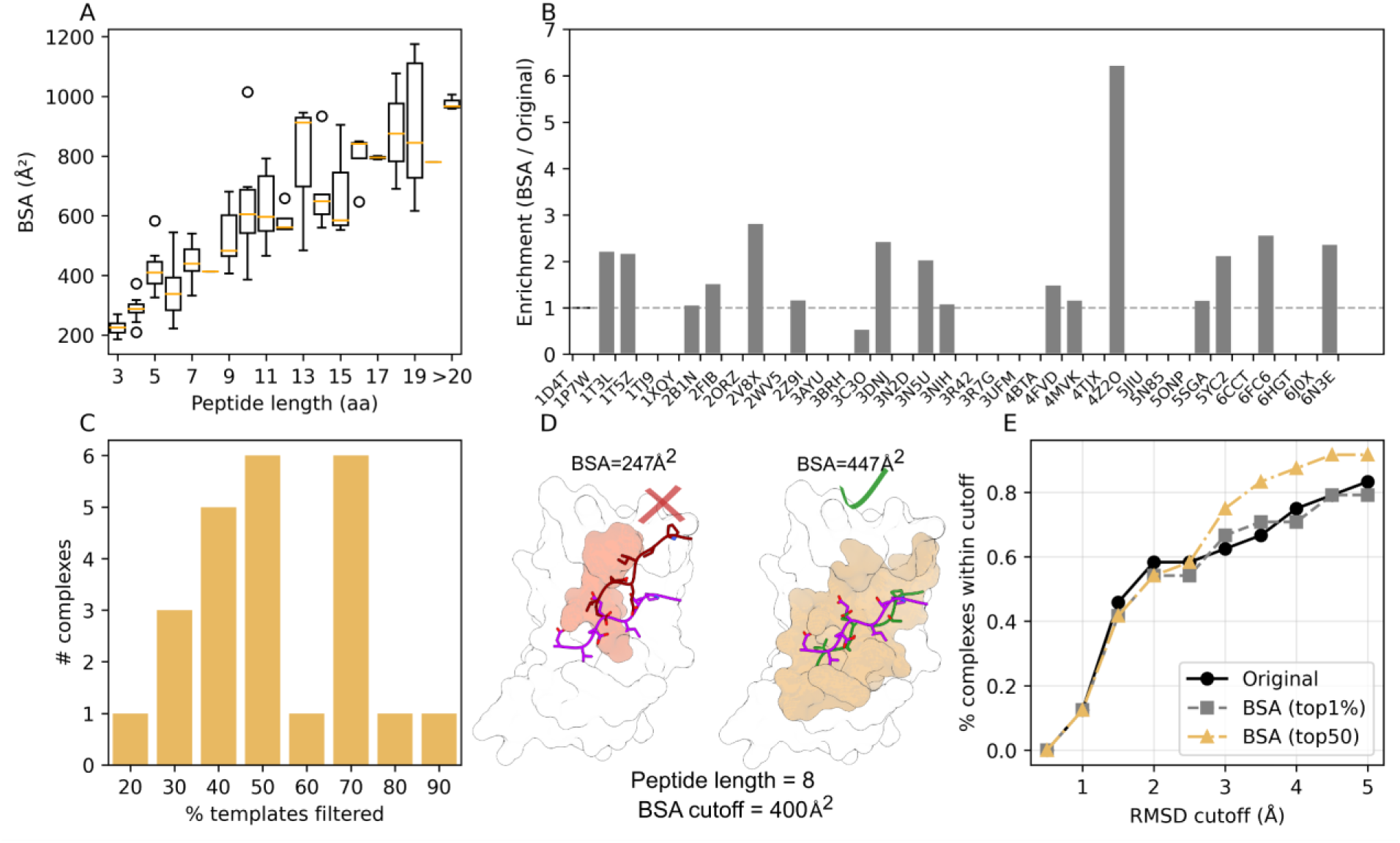
Filtering based on BSA significantly reduces the number of fragments without affecting docking performance. **A**. BSA scales with peptide length. Ranges of BSA values are shown for different peptide lengths (LNR set^29^; n=96 complexes; boxplots covering middle quartiles, divided by a line for the medians). This distribution was used to determine BSA thresholds (see **Supplementary Table S2)**. **B**. BSA filtering enrichment for good fragments (RMSD ≤2.5 Å among the top1% models) in the LNR set. **C.** Histogram of percentage of templates filtered out per complex by implementing the BSA filter on the LNR set. Values on the x-axis represent the upper limit of each bin. **D.** Example of acceptable (right) and unacceptable (left) peptide fragment conformation, based on the BSA filter (PDB 1ELW). The native peptide is colored magenta on both structures, and buried surface areas are shown in orange-brown colors. **E**. Summary of docking performance on the PFPD set^25^ (n=24 complexes), shown as the fraction of complexes (y-axis) that are modeled within a given RMSD cutoff (x-axis). (PatchMAN original: black circles; PatchMAN2 BSA clustering of top 1% scored models: gray squares; PatchMAN2 BSA clustering of minimum top50 models: yellow triangles). BSA (top50) is used as the baseline for the following analyses.

**Figure 2.**
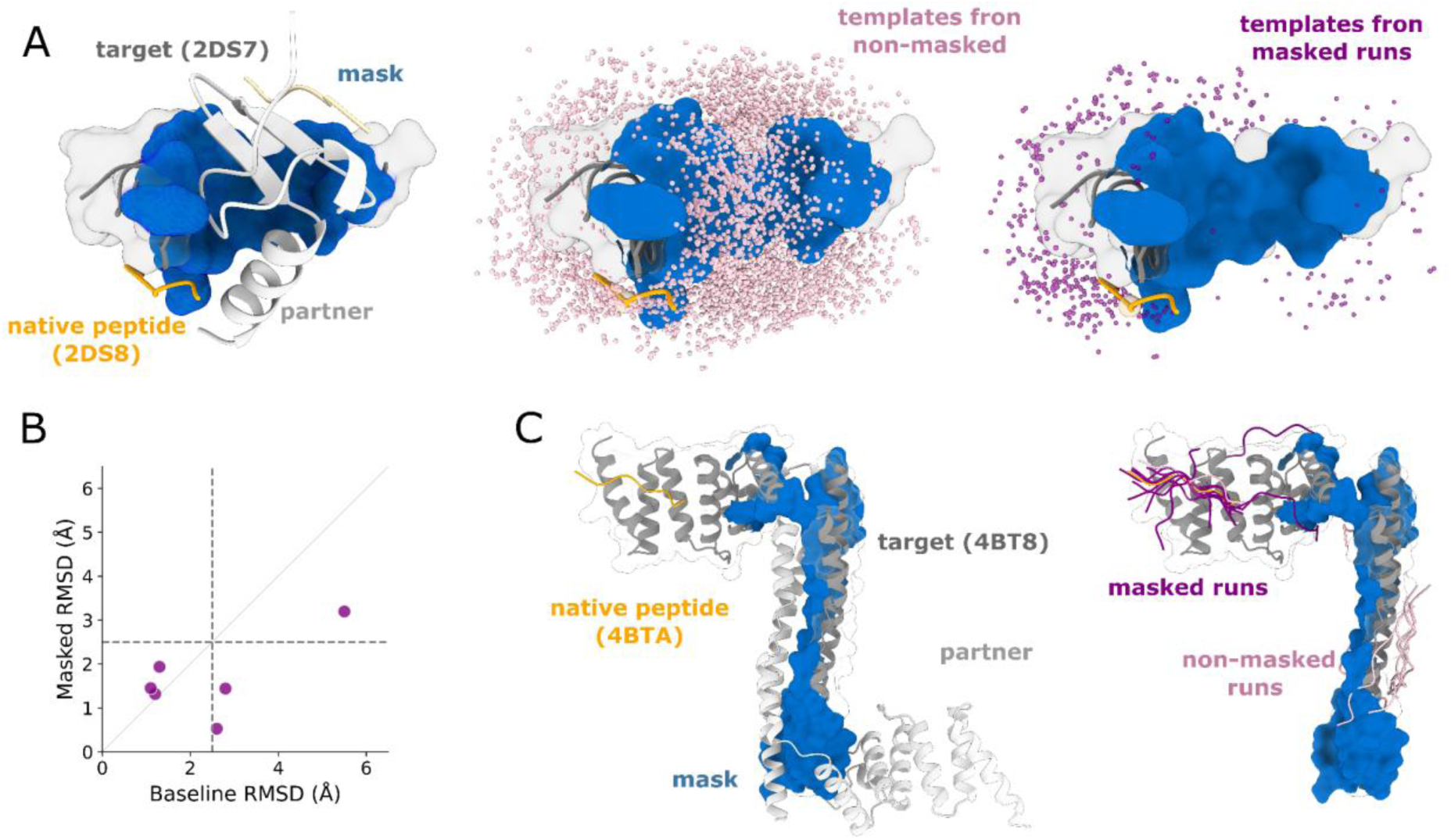
Effect of masking protein-protein interfaces on fragment extraction and docking. **A.** Overview of masking. Protein-protein interface residues used to define the mask are shown in blue on the receptor surface (white), with the peptide shown as an orange cartoon. Fragment positions obtained without masking (**middle panel**) and after masking (**right panel**) are shown in pink and purple, respectively. **B.** Docking performance before and after masking, reported as RMSD of the best model among the top 10 cluster representatives. **C.** Structural example for complex 4BTA. The masked region is shown in blue and the binding partner in light gray (**left**). Top 10 models obtained before and after masking are shown in pink and purple, respectively (**right**).

**Figure 3.**
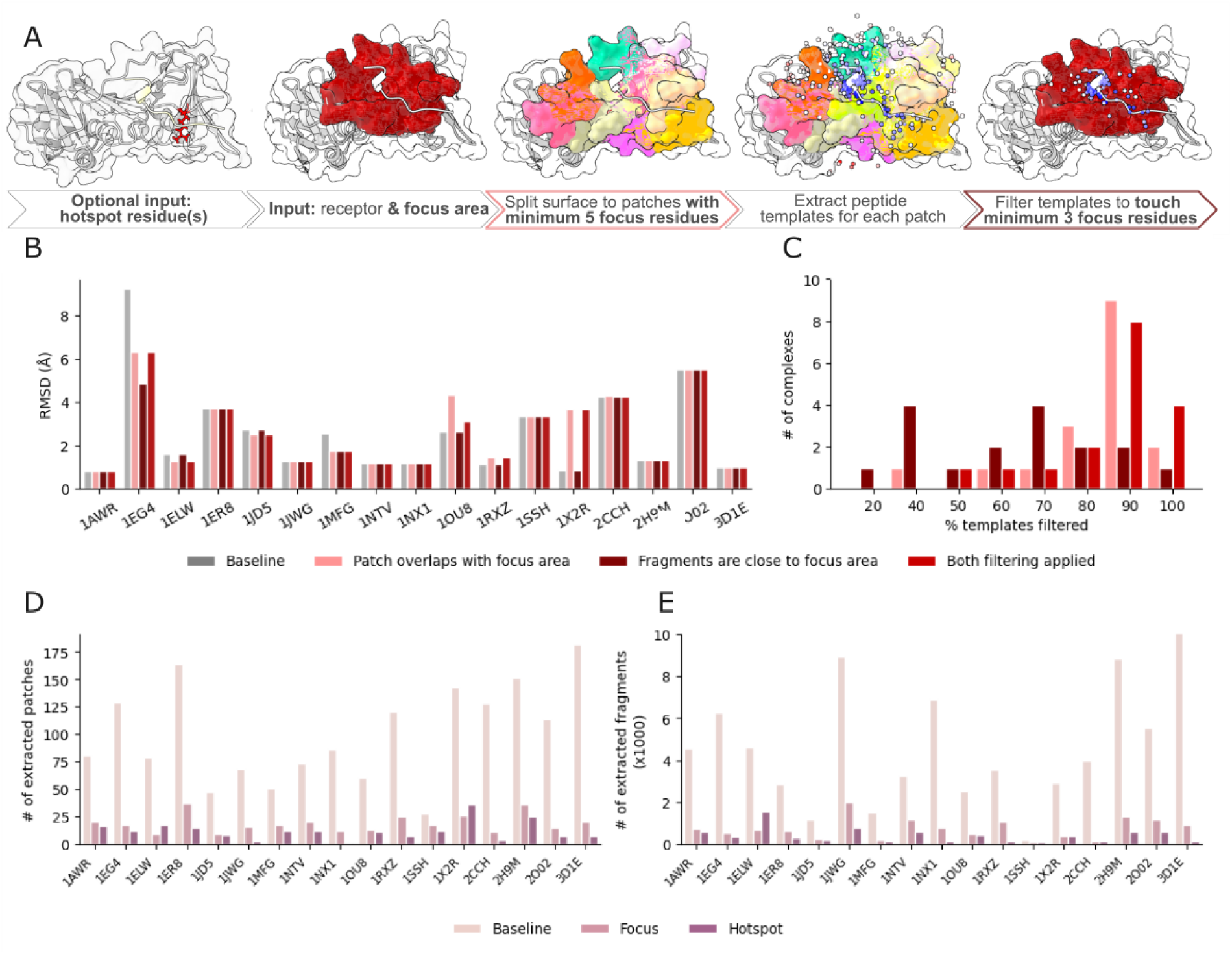
Calibration of PatchMAN2 focus and hotspot protocols on the PFPD benchmark set. **A.** Overview of the PatchMAN2 focus and hotspot workflow. The focus area and hotspot residues on the unbound receptor (PCNA; PDB ID 1RWZ) are shown in red, and extracted patches are displayed in different colors. Dots represent C_α_ atoms of fragment center residues prior to refinement, colored by RMSD to the native structure after refinement (blue, low RMSD; red, high RMSD). The native peptide (PDB ID 1RXZ) is shown in white. **B.** Effect of focus filtering on docking performance (after clustering of refined models). **C.** Effect on fragment selection: Percentage of templates filtered (as in Fig. 1C). **D-E.** Reduction in the number of extracted patches (**D**) and fragments (**E**) following focus and hotspot filtering.

### 3.1. BSA filtering during template extraction

The buried surface area (BSA) of protein-protein complexes correlates with binding affinity^40^ and is commonly used to assess model quality. However, in peptide-protein complexes the total BSA is smaller and strongly depends on peptide conformation. To evaluate whether BSA can be used to filter unpromising peptide template fragments, we calculated BSA values for native peptide-protein complexes from the LNR dataset^29^ (n = 96; Methods), which includes helical, β-strand, and coiled peptide conformations. Because BSA scales with peptide length, we derived length-dependent thresholds from the distribution of native complexes (**Fig. 1A; Supplementary Table S2**; Methods).

Overall, applying BSA filtering enriched fragments that produce near-native successful models (RMSD ≤ 2.5 Å, calculated over peptide interface backbone atoms, as rmsBB_if in FPD, Methods) among the top 1% scoring models for 9 of the 17 cases where such fragments were present, while removing them only in one case (3C3O) (**Fig. 1B**). As expected, BSA filtering does not significantly affect performance (**Supplementary Figure 1; Supplementary Table S3B**; measured as the best RMSD among the top 10 cluster representatives after re-clustering the top-scoring models; Methods), while substantially reducing the number of fragments retained and the resulting computational cost (**Fig. 1C**).

Because BSA-based filtering reduces the number of retained fragments, clustering using only the top 1% of models (as in the original PatchMAN implementation) may insufficiently capture the conformational space sampled for small systems. To avoid clustering artifacts when few fragments remain after filtering, we enforced a minimum of 50 models for clustering (*max[top50, top1%]*, hereafter referred to as **top50**). This did not change performance (**Supplementary Figure 1; Supplementary Table S3B**).

To confirm that BSA filtering does not compromise docking accuracy, we applied the derived cutoffs to the results of the original PatchMAN application on the independent PFPD set^25,27^ (n=24), which was not used for the above cutoff derivation. Template complexes were filtered prior to refinement using the defined cutoffs (**Supplementary Table S2). Fig. 1D** shows an example for excluded and included 8-residue peptide fragments. Filtering did not significantly affect high resolution docking performance (range within 2.5 Å RMSD) but slightly improved medium resolution performance (3-5 Å RMSD range) (**Fig. 1E**). One exception was PDB 1SSH, where all fragments producing near-native models were removed, increasing the final RMSD from 1.3 Å to 3.3 Å (**Supplementary Table S3A**). In the following we refer to the PatchMAN2 results obtained with BSA filtering and top50 clustering as **PatchMAN2_baseline_**.

### 3.2. Masking irrelevant receptor surfaces

Sampling can be reduced by excluding receptor surface regions unlikely to participate in peptide binding, for example obligatory protein-protein interfaces in multimeric complexes. To enable this, we introduced an option to define *masked regions* on the receptor surface (**Fig. 2A**).

Masking is applied at two stages of the PatchMAN workflow. First, during patch generation, surface patches that substantially overlap with the masked region are removed (N_mask_overlap_ ≥ 30% of patch residues). Second, during template extraction from MASTER hits, fragments that contact multiple masked residues are discarded (N_mask_contacting_ ≥ 3 residues).

To evaluate the effect of masking, we analyzed homomultimeric complexes whose obligatory interfaces had been manually masked in the original PatchMAN study^27^ (1NX1, 2DS8, 1OU8, 2O02, 1CZY, 4BTA, 1RXZ; **Supplementary Table S4**). These examples illustrate how strongly such interfaces can bias fragment extraction in the absence of masking (**Fig. 2A**). Masking only ∼10-15% of the receptor surface reduced the number of extracted fragments by ∼30%, while masking ∼30% of the surface removed up to 70-90% of fragments in some cases (**Supplementary Figures S2-S3**).

Importantly, masking misleading interfaces decreases runtime and can improve docking performance by preventing fragments from clustering at non-relevant interaction sites (**Fig. 2B**). For example, in 4BTA, masking the protein-protein interface redirected sampling toward the native peptide-binding region: after masking, the top-ranked models converged near the correct binding site, whereas without masking most models overlapped with the masked interface (**Fig. 2C**).

### 3.3. Guided docking using focus regions

To further improve speed and performance, we investigated ways to optimally guide docking toward relevant regions of the receptor surface. We therefore introduced a filtering option that restricts sampling to a defined focus area on the receptor (**Fig. 3A**). Such regions can be defined using experimental information, such as homologous complex structures, or predicted binding sites obtained by different approaches^12–19^. In addition, we describe the option of hotspot-based inputs in Section 3.4.

#### 3.3.1. Calibration of PatchMAN2 focus filtering mode

To calibrate the focus filtering parameters, we reanalyzed the PatchMAN2_baseline_ runs using homologous complex structures to define candidate binding regions. Suitable homologs were identified for 17 of the 24 complexes in the PFPD benchmark set (**Supplementary Table S1; Methods**), of which 10 contained promising fragments and were therefore used for calibration.

We analyzed filtering in two steps of the PatchMAN pipeline: the surface patch selection step and the template extraction step from MASTER hits (**Fig. 3A**, see also Methods): Successful patches typically overlapped with at least six focus residues (**Supplementary Fig. S4A**), leading us to select a threshold of N*_focus_overlap_* ≥ 5 to retain relevant patches while avoiding overfitting. Similarly, successful fragments extracted from MASTER hits consistently contacted at least three focus residues (**Supplementary Fig. S4B**), and we therefore set N*_receptor_focus_contacts_* ≥ 3. These filters increased the fraction of fragments producing near-native successful models and improved the proportion of such models among the top-ranked predictions (**Supplementary Fig. S4C and S4D**). Clustering only the top-ranking models obtained from the refinement of retained fragments resulted overall in similar (or slightly better) performance (**Fig. 3B**), while substantially decreasing the number of fragments subjected to refinement (**Fig. 3C-F**).

#### 3.3.2. Recalibration of local refinement for focused docking

Because focus filtering dramatically reduces the number of candidate fragments entering the refinement stage (**Fig. 3E**), it enables deeper sampling of relevant conformations while maintaining manageable computational cost. We therefore evaluated refinement parameters for this setting. Generating multiple models per fragment improved conformational sampling while reducing stochastic variability (**Supplementary Fig. S5**). Performance improvements plateaued beyond three refinement runs per fragment, and we therefore set *nstruct = 3*. As for BSA filtering, selecting at least 50 models (*max[top50, top1%]*) provided robust clustering across runs with varying numbers of generated models **(Supplementary Fig. S5)**. These settings were therefore adopted for subsequent focused docking runs.

#### 3.3.3. Implementation of calibrated focus mode on the benchmark datasets

Applying the calibrated focus protocol to the PFPD benchmark set improved docking success (within 2.5 Å RMSD) from 9/17 complexes using the global PatchMAN2_baseline_ to 13/17 complexes, while requiring refinement of only a fraction of the fragments (**Fig. 4A**; **Supplementary Table S6**). Notably, all but one complex (16/17) were modeled within 5.0 Å RMSD. Representative examples illustrate the impact of focus filtering on sampling and docking accuracy. For 1ER8^41^ (**Fig. 4B, left**), restricting sampling to the focus area improved both runtime and docking accuracy by directing sampling toward relevant conformations. In contrast, 1EG4^42^ (**Fig. 4B, middle**) illustrates a limitation of the approach: the focus region derived from a homologous structure does not overlap well with the native binding site, resulting in misleading patches and poor sampling. For 1JD5^43^ (**Fig. 4B, right)**, performance decreased due to differences in the peptide flanking regions between the homolog and native complexes. Notably, for 1X2R, re-running the protocol improved the final RMSD from 3.7 Å to 0.7 Å (**Supplementary Tables S5-S6**), likely due to the increased number of refinement runs enabled by the focused protocol (**Supplementary Fig. S5**).

**Figure 4.**
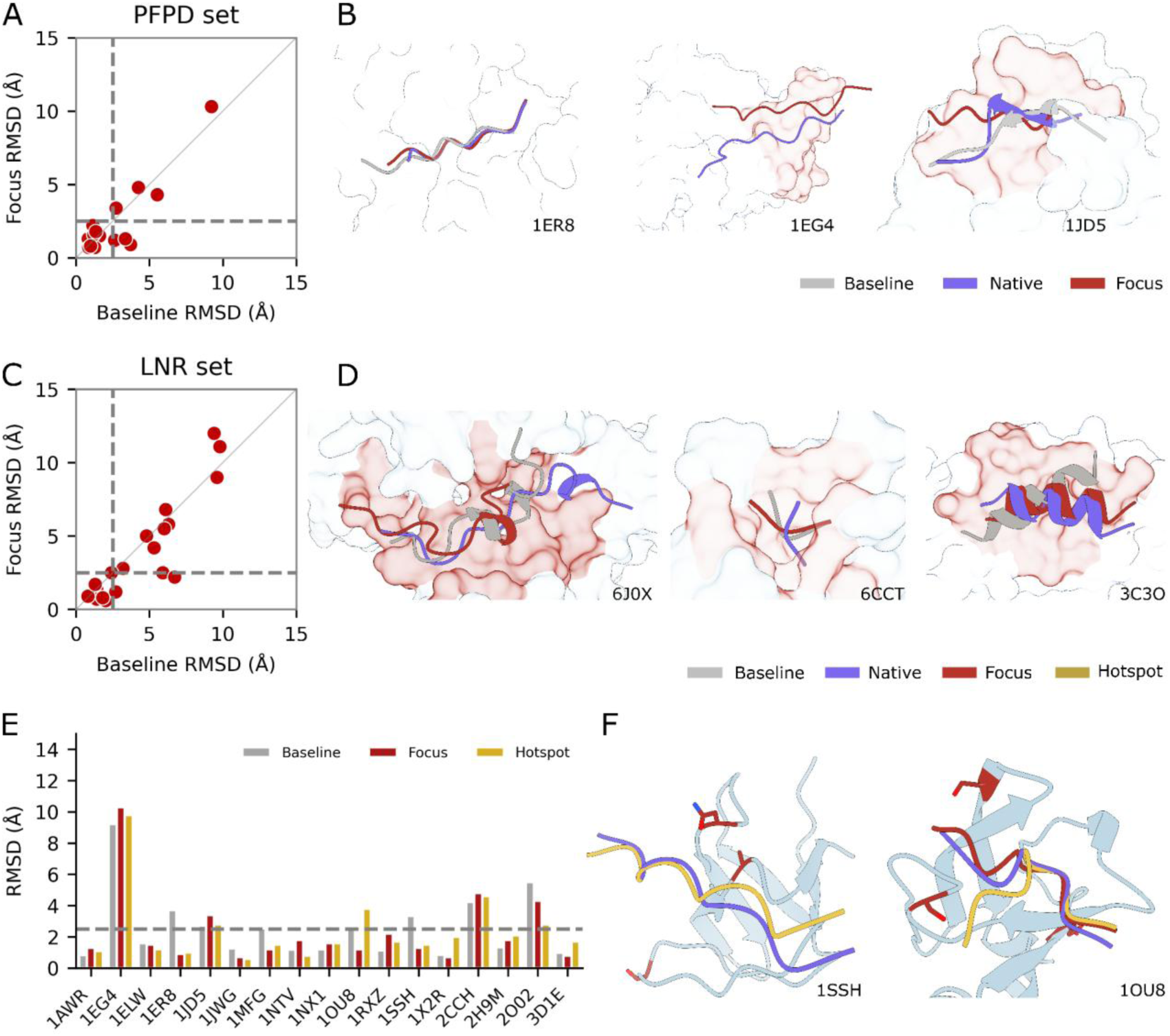
Effect of focus and hotspot filtering on docking performance compared to PatchMAN2_baseline_. **A-D.** Focus filtering results. **A.** Scatter plot comparing docking performance obtained with PatchMAN2_baseline_ and focus-guided PatchMAN2 on the PFPD benchmark set. Each point represents one complex; performance is reported as the best RMSD among the top 10 cluster representatives. The dashed line indicates equal performance between the two protocols. **B.** Structural examples illustrating the effect of focus filtering. Native peptide structures are shown together with models generated by PatchMAN2_baseline_ and focus runs for complexes 1ER8, 1EG4, and 1JD5 (left to right). **C.** As in **A.,** but for the LNR validation set. **D.** Structural examples illustrating different outcomes of focus filtering for complexes 6J0X, 6CCT, and 3C3O (left to right). **E-F.** Hotspot filtering results. **E.** Comparison of docking performance obtained using hotspot filtering, focus filtering, and PatchMAN2_baseline_ on the PFPD benchmark set. **F.** Structural examples of hotspot-guided docking for complexes 1SSH and 1OU8.

To further validate the protocol, we applied the calibrated focus pipeline to the independent LNR benchmark set. Focused docking generated successful models for 12/21 complexes, a similar or improved performance compared to PatchMAN2_baseline_ (**Fig. 4C**; **Supplementary Table S6B**). For example, in 3C3O (**Fig. 4D, right**), a well-defined focus region effectively guided sampling toward the correct binding site and substantially improved performance. In seven cases, no models within 5 Å RMSD were selected (**Supplementary Table S6B**). These failures were primarily due to insufficient sampling (e.g., 6J0X^44^, **Fig. 4D left**) or scoring limitations, where near-native conformations were sampled but ranked poorly (e.g., 6CCT^45^, **Fig. 4D center**, best RMSD 1.9 Å ranked > 500).

Overall, the protocol performs robustly on the LNR set, with results comparable to those observed for the PFPD dataset; remaining failures are mainly due to either imperfectly defined focus areas or limitations of the Rosetta scoring function, which sometimes fails to rank high-quality models among the top predictions.

In summary, focus filtering reduces the number of fragments entering the refinement stage and therefore lowers computational cost while maintaining, and in some cases improving, docking accuracy. This strategy is particularly effective when the binding site is known or can be reliably predicted, and for receptors with extended or multi-surface interaction regions.

### 3.4. Docking guided by binding hotspot residues

Besides defining broader focus regions, in many experimental situations only a small number of key interface residues are known, for example from mutational scanning experiments. We therefore investigated whether PatchMAN2 can use binding hotspot residues as minimal input to guide docking. To simulate such cases, we performed computational alanine scanning on the native complexes and selected the three residues with the largest predicted contribution to binding energy (ΔΔG_binding_) as hotspots (**Supplementary Table S1**; Methods). The hotspot region was then expanded to include nearby residues to define a local interaction area for filtering (**Fig. 3A**).

Because hotspot regions are typically smaller than focus areas (**Supplementary Fig. S6**), we relaxed the template filtering threshold to require only one contacting hotspot residue (N*_receptor_hotspot_contacts_* ≥ 1). As with focus filtering, hotspot filtering substantially reduced the number of patches and fragments retained for refinement (**Fig. 3D-E**), thereby decreasing runtime.

#### 3.4.1. Application of PatchMAN2 hotspot filtering mode to the PFPD benchmark set

Applying hotspot-guided docking to the PFPD benchmark set affected performance in several cases (**Fig. 4E**). For example, in 1SSH, hotspot filtering improved modeling compared to the global PatchMAN2_baseline_ (**Fig. 4F, left; Supplementary Table S6A**). In contrast, for 1OU8 hotspot filtering did not reduce the final RMSD relative to the baseline or focus runs (**Fig. 4F, right; Supplementary Table S6A**). In this case, near-native conformations were sampled (∼3 Å RMSD), but were not selected among the final models due to ranking and clustering. Overall, these results demonstrate that useful docking predictions can be obtained even with very limited information about the interaction region. However, as in the focus protocol, successful hotspot-guided docking depends on accurate identification of the hotspot residues.

## Discussion

Peptide docking remains challenging because peptides often adopt a defined structure only upon binding. This coupling between folding and binding greatly expands the conformational search space, as modeling must simultaneously identify both the peptide conformation and its binding mode. PatchMAN decouples these two steps by using peptide template fragments that complement receptor surface-similar structural submotifs in solved structures as starting points, that are subsequently refined using the Rosetta FlexPepDock protocol^28^. Because this refinement step involves extensive conformational sampling, reducing the number of fragments subjected to refinement is critical for improving efficiency.

Here we present PatchMAN2, a new version that includes a number of ways to accelerate PatchMAN docking by prioritizing relevant starting structures for refinement. The main goal is to remove irrelevant fragments before refinement while preserving docking accuracy. This can accelerate prediction time, or in turn, allow resources to be reallocated toward increased sampling starting from the remaining relevant fragments.

To achieve this, we first remove fragments that make only minimal contact with the receptor, and therefore are most probably not good starting points for the generation of successful models of a peptide-receptor complex. This is achieved with a peptide length-dependent threshold of buried surface area (BSA, **Fig. 1**). Furthermore, we also enable PatchMAN to mask inaccessible surfaces on the protein receptor that are occupied by obligatory binding to other proteins as homo- or hetero-multimers (**Fig. 2**). These surfaces are often sticky and will attract many false positive fragments. Moreover, we implemented a focus mode, for the frequent scenario where information is available about the binding site (for example from a solved homolog complex, or experimentally or computationally identified hotspot residues), enabling PatchMAN to concentrate on the specific surface area, as well as to filter out fragments outside of this area (**Figs. 3&4**). On established benchmarks of peptide-protein complex structures used in our previous studies we describe here a quantitative assessment of the performance of PatchMAN2 and highlight the improvement in runtime while maintaining, and often improving, performance. Results clearly demonstrate that including information about the binding site as input for PatchMAN saves runtime and increases the chance of success. Importantly, these filters allow PatchMAN2 to incorporate biological knowledge directly into the docking process, enabling users to guide sampling using structural constraints, predicted binding regions, or sparse experimental information such as hotspot residues.

Conceptually, PatchMAN local docking resembles ab-initio folding of the peptide into the binding pocket. While most local docking methods rely on extracting and refining peptide templates from already solved, homologous peptide-protein complexes, this often cannot move the backbone enough to sample vastly different conformations, possibly missing novel conformations that have not been solved previously. To tackle this problem, we previously developed Rosetta FlexPepDock *ab-initio,* which folds peptides within a binding pocket, but with moderate success and high resource usage^46^. The focused PatchMAN2 protocol presented in this study extracts peptide templates from many different structures, inherently localizing sampling to relevant conformations from the start, making it more efficient and accessible.

Across all strategies implemented in PatchMAN2, we report a consistent outcome: filtering reduced runtime significantly while maintaining - or even improving - the quality of results and performance. These enhancements extend PatchMAN’s utility across diverse docking scenarios, enabling its application in both exploratory modeling and targeted, information-driven predictions. To ensure applicability, code and setup instructions can be found at https://github.com/Furman-Lab/PatchMAN/, and a dedicated server is also available at https://r2.graylab.jhu.edu/apps/submit/patchman/.

This study also has its limitations: while we prioritized discussing run time, increasing sampling is often beneficial when computationally feasible, as it enables better coverage of the conformational space. However, despite improved sampling, scoring and ranking the generated models remain a major challenge. In particular, Monte Carlo sampling of FPD might sometimes result in different outcomes, and the currently used Rosetta score^47^ (specifically, the reweighted score that up-weighs the peptide and peptide-receptor interface energy) can miss high-quality models or rank them poorly. Improved ranking strategies - potentially incorporating machine learning, interface-specific scoring, or consensus methods - will further enhance the reliability of final model selection.

In the AI era, much of the modeling efforts have been significantly simplified by deep learning models such as Alphafold2, Alphafold3, Chai-1 and RoseTTAFold-All-Atom, HelixFold3 and more^29,48^. While these methods perform well when similar examples exist in their training data, it remains unclear how well they extend to truly novel interactions without resolved templates or similar representations^11^. Additionally, even AI-based methods struggle to confidently identify which structural solution most closely resembles the native conformation using available scoring functions^49^. Physics-based and motif-based approaches such as PatchMAN therefore provide an important complementary strategy, particularly when structural motifs rather than sequence similarity determine binding. Combining such approaches with modern AI-based predictions may provide the most robust framework for modeling peptide-protein interactions across diverse biological scenarios.

## Code and data availability

The pipeline is available for local running at https://github.com/Furman-Lab/PatchMAN.

## Supporting information

The datasets, the databases used for MASTER search and extraction of fragments, and the files containing the PDB id-s excluded from the search for benchmarking have been uploaded to Zenodo (https://doi.org/10.5281/zenodo.16567267).

## Acknowledgements

This work was partly supported by the Israel Science Foundation (ISF) founded by the Israel Academy of Sciences and Humanities (grant #s 301/2021 and 3091/23), and by the European Union Horizon 2020 Research and Innovation program under the Marie Skłodowska-Curie Grant Agreement (UBIMOTIF; grant # 860517).

