## Supplementary for "Strategic template filtering accelerates fragment-based peptide docking"

### Supplementary Information

#### Supplementary Figures

- **Supplementary Figure S1**: Performance for the LNR set (n=39): comparison of the original PatchMAN results with PatchMAN2 which includes BSA filtering and clustering of decoys using either the top 1% [BSA(top 1%)] or max(top 50, top 1%) [BSA(top50)]. BSA(top5) is the baseline for the following analysis ($PatchMAN_{baseline}$). Legend as in **Figure 1E.**
- **Supplementary Figure S2:** Masking interfaces of obligatory homomultimers
- **Supplementary Figure S3:** Masking reduces the number of extracted patches and fragments
- **Supplementary Figure S4**: Calibration of filtering thresholds on the PFPD set.
- **Supplementary Figure S5**: Calibration of number of refinement runs and decoys selected for clustering.
- **Supplementary Figure S6:** Comparison of receptor surface area covered by hotspot-defined input regions and focus-defined regions.

#### Supplementary Tables

- **Supplementary Table S1**: The two benchmark sets used in this study, composed of complexes for which a suitable bound homologue structure is available (with a peptide located within 6 Å RMSD of the peptide in the benchmark structure).
- **Supplementary Table S2**: Definition of BSA cutoff for different peptide lengths
- **Supplementary Table S3:** Performance of PatchMAN after addition of BSA filter
- **Supplementary Table S4**: Masking the obligatory interface of homomultimer complexes improves PatchMAN results
- **Supplementary Table S5**: Effect of focus filters on performance (PFPD set)
- **Supplementary Table S6**: Performance of PatchMAN after addition of focus filtering

#### Supplementary Files

- **Supplementary File S1:** List of PDBs with the same receptor that are excluded in the MASTER search step

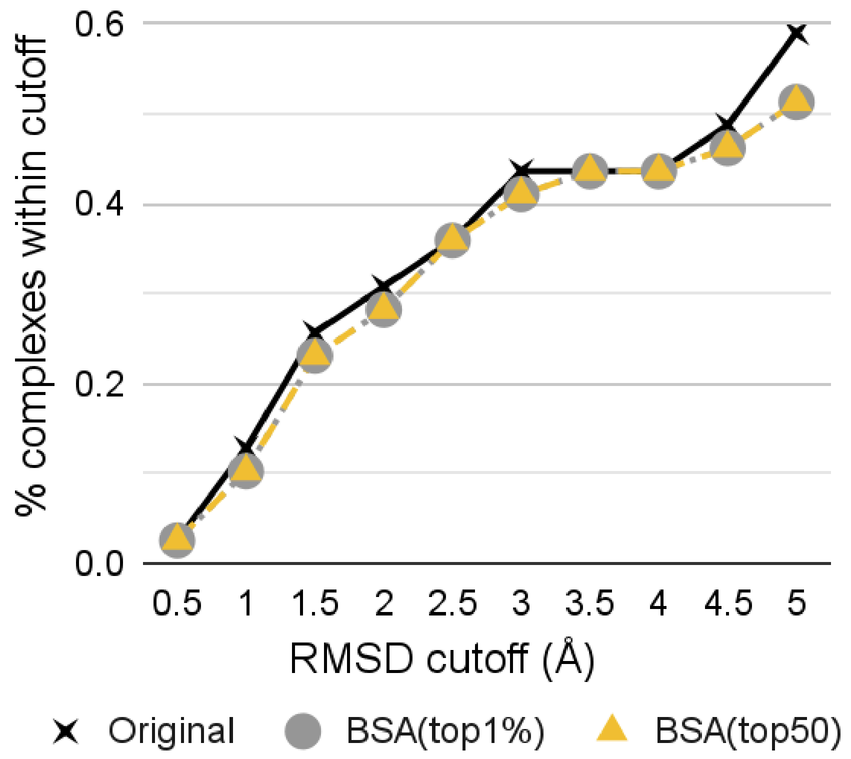


**Supplementary Figure S1.** Performance for the LNR set (n=39): Comparison of the original PatchMAN results with PatchMAN2 which includes BSA filtering and clustering of decoys using either the top 1% [BSA(top1%)], or *max(top 50, top1%)* [BSA(top50)]. BSA(top50) is the baseline for the following analysis ($\text{PatchMAN}_{\text{baseline}}$). Legend as in **Figure 1E.**

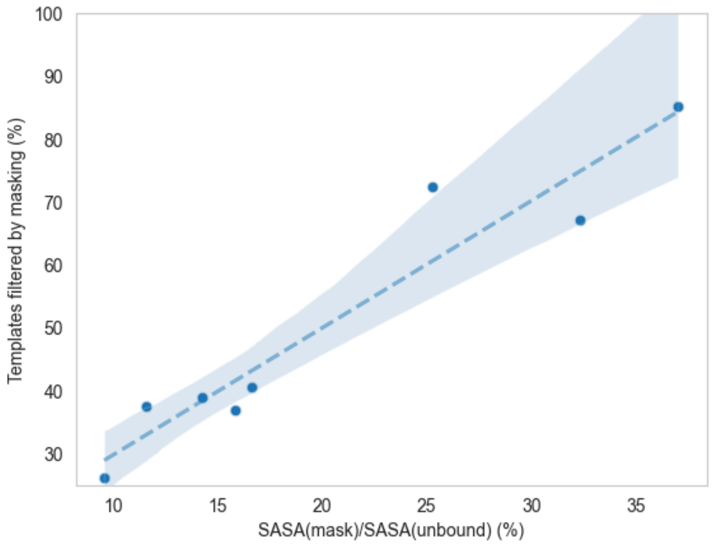


**Supplementary Figure S2.** Masking interfaces of obligatory homomultimers. Masking one third of the surface can result in the removal of 80-90% of the candidate fragments, due to the popularity of the interface for binding. The percentage of fragments filtered by masking is correlated with the percentage of the receptor surface covered by the multimer interface. SASA - Solvent Accessible Surface Area.

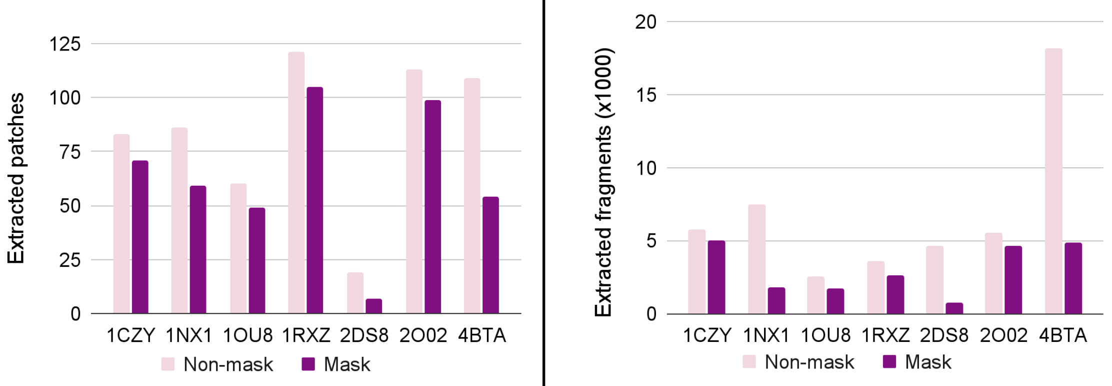


**Supplementary Figure S3.** Masking reduces the number of extracted patches (**left**) and fragments (**right**).

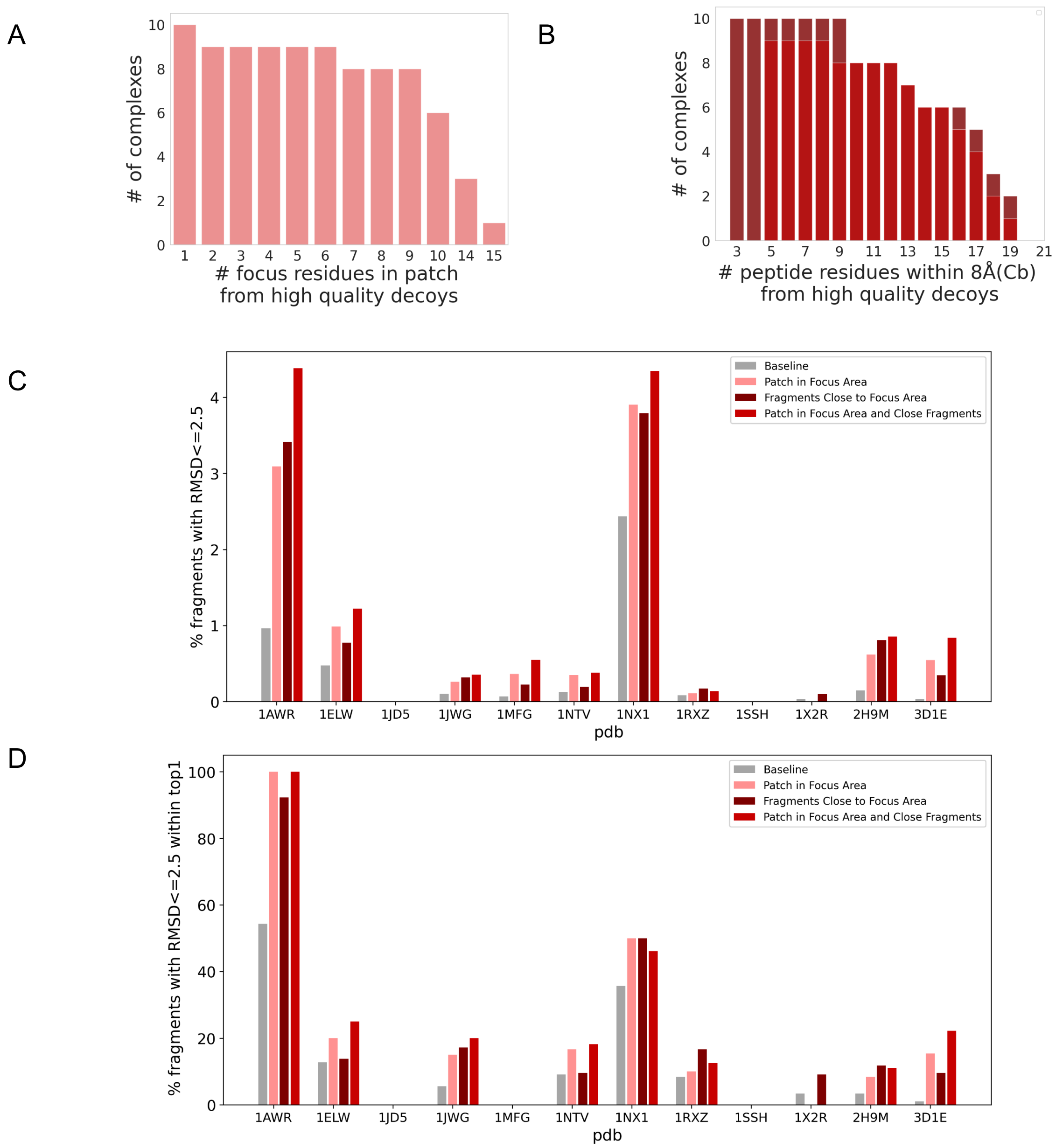


**Supplementary Figure S4.** Calibration of filtering thresholds on the PFPD set. **A.** Distribution of the number of overlapping residues between the focus area and patches producing successful models. **B**.Distribution of the number of focus residues contacting fragments that produce successful models**. C-D.**Effect of focus filtering on fragment selection . Filtering enhances the selection efficiency of accurate models within the top-ranked predictions for the PFPD benchmark set. **C.** Fraction of fragments producing near-native models(RMSD within 2.5 Å after their refinement;). **D.** Fraction of fragments that lead to final successful models (RMSD within 2.5 Å after their refinement; among the top 1% models retained for the final clustering step in PatchMAN).

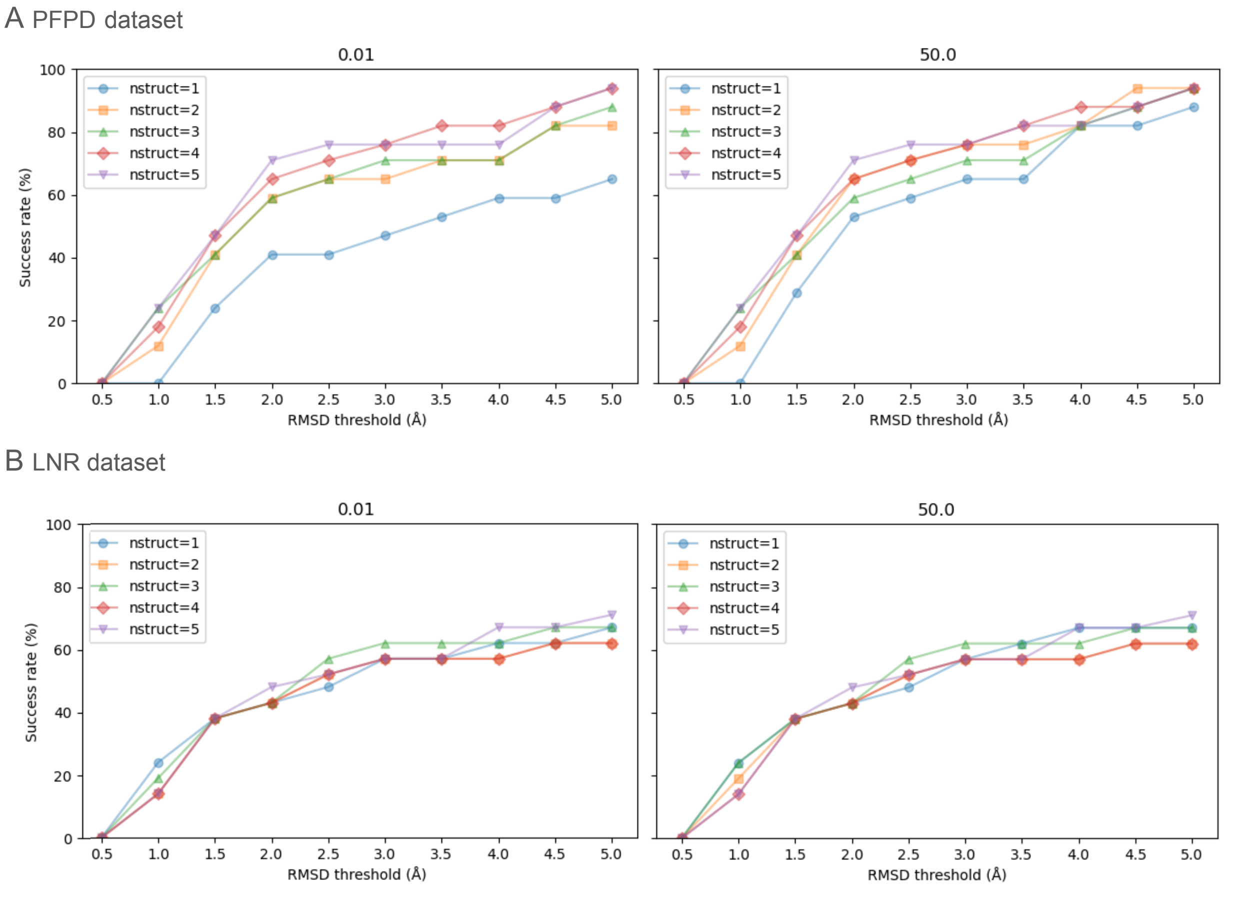


**Supplementary Figure S5.** Calibration of number of refinement runs and decoys selected for clustering for focus runs. Results are shown for benchmark sets PFPD **(A)** and LNR (**B**). Decoys are selected for clustering according to a fixed fraction of the total decoy set (**right panels**), or a fixed number of decoys (**left panels**). One to five refinements per fragment are evaluated (see color code inset). The most significant improvement is observed for increasing *nstruct,* in particular increasing to *nstruct >1*, and for some further increasing to *nstruct >3*.

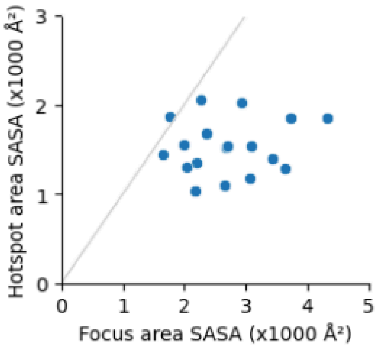


**Supplementary Figure S6.** Comparison of receptor surface area covered by hotspot-defined input regions (after expansion; see Methods) and focus-defined regions. The gray line indicates equal surface area coverage.

**Supplementary Table S1**. The two benchmark sets used in this study, composed of complexes for which a suitable bound homologue structure is available (with a peptide located within 6 Å RMSD of the peptide in the benchmark structure).

**A.** PFPD benchmark set (17/24 complexes with available homologue structure).

| Complex pdb | Unbound pdb | Homologue for focus | Focus size (# residues) | Hotspot residues |
|---|---|---|---|---|
| 1AWR:CI | 2ALF | 1CWB:A | 25 | 62,54,59 |
| 1EG4:AP | 1EG3 | 1F4V:A | 28 | 24,131,13 |
| 1ELW:AC | 1A17 | 1H27:B | 14 | 83,49,18 |
| 1ER8:EI | 4APE | 1NX0:B | 44 | 79,222,219 |
| 1JD5:AB | 1JD4 | 1OEB:A | 15 | 72,68,73 |
| 1JWG:BD | 1JWF | 2QAS:A | 20 | 86,96,82 |
| 1MFG:AB | 2H3L | 2QT5:B | 20 | 34,27,83 |
| 1NTV:AB | 1P3R | 3G2S:A | 26 | 91,134,108 |
| 1NX1:AC | 1ALV | 3UQR:A | 26 | 73,35,80 |
| 1OU8:BD | 1OU9 | 3UVK:A | 22 | 28,75,53 |
| 1RXZ:AB | 1RWZ | 3UW5:B | 34 | 241,44,242 |
| 1SSH:AB | 1OOT | 4CGW:B | 22 | 38,54,17 |
| 1X2R:AB | 1X2J | 4TSZ:A | 21 | 92,160,249 |
| 2CCH:DF | 1H1R | 5NJK:D | 19 | 80,106,50 |
| 2H9M:CD | 2H14 | 6EJL:B | 37 | 229,18,102 |
| 2O02:AP | 2BQ0 | 6K3A:C | 31 | 124,173,42 |
| 3D1E:AP | 3D1G | 6KKG:A | 28 | 175,247,362 |

**B.** LNR benchmark set (21/96 complexes with available homologue structure).

| Complex pdb | Unbound pdb | Homologue for focus | Focus size (# residues) |
|---|---|---|---|
| 1D4T:AB | 1D1Z:A | 1LKL:A | 27 |
| 1T3L:AB | 1T3S:A | 1T0J:B | 32 |
| 1T5Z:AB | 1E3G:A | 6KTM:A | 27 |
| 1TJ9:AB | 2QU9:A | 1JQ9:A | 29 |
| 2B1N:AB | 2B1M:A | 1MHW:B | 14 |
| 2FIB:AB | 3FIB:A | 1FZF:F | 21 |
| 2V8X:AB | 5GW6:A | 5EKV:A | 24 |

| | | | |
|---|---|---|---|
| 2WV5:CG | 2J92:A | 3SJ9:A | 31 |
| 2Z9I:AG | 1Y8T:A | 7CO5:F | 40 |
| 3AYU:AB | 1QIB:A | 4JQG:B | 30 |
| 3C3O:AB | 2OEW:A | 5MK3:A | 26 |
| 3N2D:AB | 3RL9:A | 2JDL:A | 29 |
| 3N5U:AC | 6G0J:A | 2P6B:A | 20 |
| 4BTA:AC | 4BT8:A | 2V5F:A | 19 |
| 4Z2O:AP | 4Z27:A | 1KL3:A | 32 |
| 5N85:AB | 4IPG:A | 5EAY:B | 27 |
| 5YC2:AB | 5YBX:A | 5YCA:A | 23 |
| 6CCT:AB | 6CCR:A | 6WZX:A | 19 |
| 6HGT:DH | 2GP5:A | 4HON:A | 42 |
| 6J0X:AE | 6J0V:A | 6J0W:A | 52 |
| 6N3E:AB | 6N3F:A | 2PEH:B | 21 |

**Supplementary Table S2.** Definition of BSA cutoff for different peptide lengths

| Peptide length | Minimum BSA (freeSASA*) [Å$^2$] | Cutoff (freeSASA) [Å$^2$] | Minimum BSA (pyRosetta**) [Å$^2$] | Cutoff (pyRosetta) [Å$^2$] |
|---|---|---|---|---|
| 3 | 185 | 100 | 239 | 200 |
| 4 | 210 | 200 | 279 | 200 |
| 5 | 326 | 200 | 405 | 200 |
| 6 | 222 | 200 | 291 | 200 |
| 7 | 332 | 300 | 384 | 300 |
| 8 | 412 | 300 | 473 | 400 |
| 9 | 406 | 300 | 457 | 400 |
| 10 | 386 | 300 | 541 | 500 |
| 11 | 466 | 400 | 518 | 500 |
| 12 | 554 | 400 | 599 | 500 |
| 13 | 484 | 400 | 516 | 500 |
| 14 | 560 | 500 | 610 | 600 |
| 15 | 552 | 500 | 625 | 600 |
| 16 | 647 | 600 | 717 | 600 |
| 17 | 789 | 600 | 823 | 600 |
| 18 | 690 | 600 | 721 | 600 |
| 19 | 616 | 600 | 699 | 600 |
| 20 | 780 | 700 | 830 | 800 |
| 21 | 967 | 900 | 1083 | 1000 |
| 27 | 959 | 900 | 1055 | 1000 |

***** as calculated using FreeSASA[1]
****** as calculated using PyRosetta[2]

**Supplementary Table S3. Performance of PatchMAN after addition of BSA filter.** Comparison of PatchMAN with PatchMAN2. Significantly improved and worse cases are highlighted in bold and italics, respectively.

A. PFPD set

| PDB | | PatchMAN* | | |
|---|---|---|---|---|
| Complex | Unbound | Original (PatchMAN) | PatchMAN2 (BSA Top 1%) | PatchMAN2 (BSA Top 50) ($PatchMAN2_{baseline}$) |
| 1AWR:CI | 2ALF:A | 0.9 | 0.8 | 0.8 |
| 1CZY:CE | 1CA4:A | 2.8 | 2.8 | 2.8 |
| 1EG4:AP | 1EG3:A | 19.6 | 9.2 | 9.2 |
| 1ELW:AC | 1A17:A | 2.0 | **1.6** | **1.6** |
| 1ER8:EI | 4APE:A | 9.7 | 10.3 | **3.7** |
| 1JD5:AB | 1JD4:A | 3.7 | **2.7** | **2.7** |
| 1JWG:BD | 1JWF:A | 1.4 | 1.3 | 1.3 |
| 1MFG:AB | 2H3L:A | 9.4 | 7.1 | **2.5** |
| 1NTV:AB | 1P3R:B | 1.2 | 1.2 | 1.2 |
| 1NX1:AC | 1ALV:A | 1.5 | 1.2 | 1.2 |
| 1OU8:BD | 1OU9:A | 4.2 | 4.2 | **2.6** |
| 1RXZ:AB | 1RWZ:A | 1.1 | 1.1 | 1.1 |
| 1SSH:AB | 1OOT:A | **1.3** | *5.2* | **3.3** |
| 1U00:AP | 2V7Y:A | 2.0 | 2.0 | 2.0 |
| 1X2R:AB | 1X2J:A | 0.9 | 0.9 | 0.9 |
| 2A3I:AB | 2AA2:A | 1.1 | 1.1 | 1.0 |
| 2B9H:AC | 2B9F:A | 3.4 | 3.4 | 3.4 |
| 2C3I:BA | 2J2I:B | 3.7 | **2.6** | **2.6** |
| 2CCH:DF | 1H1R:B | 4.9 | 4.2 | 4.2 |
| 2DS8:BP | 2DS7:A | 1.3 | 1.3 | 1.3 |
| 2H9M:CD | 2H14:A | 1.3 | 1.3 | 1.3 |
| 2HPL:AB | 2HPJ:A | 1.7 | 1.7 | 1.7 |

B. **LNR set.** Legend as in **A**

| PDB ID | | PatchMAN | | |
|---|---|---|---|---|
| Complex | Unbound | Original (PatchMAN) | PatchMAN2 (BSA Top1%) | PatchMAN2 (BSA Top50) ($PatchMAN2_{baseline}$) |
| 1D4T | 1D1Z | 8.3 | 9.8 | 9.8 |
| 1KY7 | 1QTS | 15 | 15.5 | 15.5 |
| 1P7W | 1EGQ | 4.2 | 4.2 | 4.2 |
| 1T3L | 1T3S | 1.4 | 1.4 | 1.4 |
| 1T5Z | 1E3G | 1.4 | 1.4 | 1.4 |
| 1TJ9 | 2QU9 | 4.9 | *6.1* | *6.1* |
| 1XQY | 1XQX | 2.9 | 2.9 | 2.9 |
| 2B1N | 2B1M | 1.4 | 1.4 | 1.4 |
| 2FIB | 3FIB | 2.1 | 2.0 | 2.0 |
| 2ORZ | 2ORX | 4.1 | *27.5* | *27.5* |
| 2V8X | 5GW6 | 0.9 | 0.8 | 0.8 |
| 2WV5 | 2J92 | 6.3 | 6.3 | 6.3 |
| 2Z9I | 1Y8T | 0.8 | 0.8 | 0.8 |
| 3AYU | 1QIB | 2.8 | 2.7 | 2.7 |
| 3BRH | 2QCJ | 8.8 | 9.9 | 9.9 |
| 3C3O | 2OEW | 4.7 | *5.9* | *5.9* |
| 3DNJ | 3G3P | 0.8 | 0.8 | 0.8 |
| 3N2D | 3RL9 | 9.6 | 9.6 | 9.6 |
| 3N5U | 6G0J | 2.9 | *3.2* | *3.2* |
| 3NIH | 3NIL | 1.0 | 1.0 | 1.0 |
| 3R42 | 3R3Q | 13.9 | 14.6 | 14.6 |
| 3R7G | 2YLF | 10.2 | 11.0 | 11.0 |
| 3UFM | 2BOO | 4.6 | 4.6 | 4.6 |
| 4BTA | 4BT8 | 3.6 | *6.7* | *6.7* |
| 4FVD | 4FVB | 1.9 | 1.9 | 1.9 |

| | | | | |
|---|---|---|---|---|
| 4MVK | 6GQZ | 2.4 | 2.4 | 2.4 |
| 4TJX | 4TJV | 16.7 | 16.7 | 16.7 |
| 4Z2O | 4Z27 | 1.7 | 1.8 | 1.8 |
| 5GR9 | 5JFK | 6.3 | 6.1 | 6.1 |
| 5JIU | 5JI9 | 7.4 | 7.4 | 7.4 |
| 5N85 | 4IPG | 7.7 | 6.0 | 6.0 |
| 5ONP | 5DLH | 21.9 | 21.9 | 21.9 |
| 5SGA | 2SGA | 0.3 | 0.3 | 0.3 |
| 5YC2 | 5YBX | 1.1 | *2.4* | *2.4* |
| 6CCT | 6CCR | 4.8 | 4.8 | 4.8 |
| 6FC6 | 6FC5 | 5.3 | 5.3 | 5.3 |
| 6HGT | 2GP5 | 5.3 | 5.3 | 5.3 |
| 6J0X | 6J0V | 15.2 | 9.4 | 9.4 |
| 6N3E | 6N3F | 1.3 | 1.3 | 1.3 |

**Supplementary Table S4**. **Masking the obligatory interface of homomultimer complexes improves PatchMAN results.** Shown are RMSD (in [Å], calculated over peptide interface residue backbone atoms) for the best prediction among representatives of the top 10 clusters.

| Complex PDB | Masked | Non-masked |
|---|---|---|
| 1CZY | **1.4** | *2.8* |
| 1NX1 | 1.3 | 1.2 |
| 1OU8 | **0.5** | *2.6* |
| 1RXZ | 1.45 | 1.1 |
| 2DS8 | 1.9 | 1.3 |
| 2O02 | **3.20** | 5.5 |
| 4BTA | **1.33** | 6.7 |

**Supplementary Table S5**. **Effect of focus filters on performance** (PFPD benchmark set). Shown are results where a bound homologue structure is available (**Supplementary Table S1**): the best prediction among representatives of the top 10 clusters (RMSD values in [Å], calculated over peptide interface residue backbone atoms).

| PDB ID | PatchMAN2$_{baseline}$ | Filter 1 $N_{focus\_overlap} \geq 5$ | Filter 2 $N_{receptor\_focus\_contacts} \geq 3$ | Focus: Filters 1&2 |
|---|---|---|---|---|
| 1AWR | 0.8 | 0.8 | 0.8 | 0.8 |
| 1EG4 | 9.2 | 6.3 | **4.9** | 6.3 |
| 1ELW | 1.6 | **1.3** | 1.6 | **1.3** |
| 1ER8 | 3.7 | 3.7 | 3.7 | 3.7 |
| 1JD5 | 2.7 | 2.5 | 2.7 | 2.5 |
| 1JWG | 1.3 | 1.3 | 1.3 | 1.3 |
| 1MFG | 2.5 | **1.8** | **1.8** | **1.8** |
| 1NTV | 1.2 | 1.2 | 1.2 | 1.2 |
| 1NX1 | 1.2 | 1.2 | 1.2 | 1.2 |
| 1OU8 | 2.6 | 4.3 | 2.6 | 3.1 |
| 1RXZ | 1.1 | 1.5 | 1.1 | 1.5 |
| 1SSH | 3.3 | 3.3 | 3.3 | 3.3 |
| 1X2R | 0.9 | *3.7* | 0.9 | *3.7* |
| 2CCH | 4.2 | 4.3 | 4.2 | 4.3 |
| 2H9M | 1.3 | 1.3 | 1.3 | 1.3 |
| 2O02 | 5.5 | 5.5 | 5.5 | 5.5 |
| 3D1E | 1.0 | 1.0 | 1.0 | 1.0 |

**Supplementary Table S6**. Results of focus and hotspot runs on complexes in the PFPD and LNR benchmark sets, where applicable (*i.e.,* the receptor had a homologue with a peptide bound, see **Supplementary Table S1**). Shown are RMSD values (in [Å], calculated over peptide interface residue backbone atoms) for the best prediction among representatives of the top 10 clusters.

**A. PFPD set**

| Complex PDB | $PatchMAN2_{baseline}$ | Focus | Hotspot |
|---|---|---|---|
| 1AWR | 0.8 | 1.3 | 1.1 |
| 1EG4 | 9.2 | 10.3 | 9.8 |
| 1ELW | 1.6 | 1.5 | 1.2 |
| 1ER8 | 3.7 | **0.9** | **1.0** |
| 1JD5 | 2.7 | *3.4* | **2.8** |
| 1JWG | 1.3 | **0.7** | **0.6** |
| 1MFG | 2.5 | **1.2** | **1.5** |
| 1NTV | 1.2 | *1.8* | **0.8** |
| 1NX1 | 1.2 | 1.6 | 1.6 |
| 1OU8 | 2.6 | **1.2** | *3.8* |
| 1RXZ | 1.1 | *2.2* | **1.7** |
| 1SSH | 3.3 | **1.3** | **1.5** |
| 1X2R | 0.9 | 0.7 | *2.0* |
| 2CCH | 4.2 | 4.8 | 4.6 |
| 2H9M | 1.3 | 1.8 | *2.1* |
| 2O02 | 5.5 | 4.3 | **2.8** |
| 3D1E | 1.0 | 0.8 | 1.7 |

**B. LNR set** Legend as in **A**

| Complex PDB | $PatchMAN2_{baseline}$ | Focus |
|---|---|---|
| 1D4T | 9.8 | 11.1 |
| 1T3L | 1.4 | 1.4 |
| 1T5Z | 1.4 | 1.4 |
| 1TJ9 | 6.1 | 6.8 |
| 2B1N | 1.4 | **0.7** |
| 2FIB | 2.0 | **0.6** |

| | | |
|---|---|---|
| 2V8X | 0.8 | 0.9 |
| 2WV5 | 6.3 | 5.8 |
| 2Z9I | 0.8 | 0.9 |
| 3AYU | 2.7 | **1.2** |
| 3C3O | 5.9 | **2.5** |
| 3N2D | 9.6 | 9.0 |
| 3N5U | 3.2 | **2.8** |
| 4BTA | 6.7 | **2.2** |
| 4Z2O | 1.8 | **0.8** |
| 5N85 | 6.0 | 5.5 |
| 5YC2 | 2.4 | 2.5 |
| 6CCT | 4.8 | 5.0 |
| 6HGT | 5.3 | 4.2 |
| 6J0X | 9.4 | *12.0* |
| 6N3E | 1.3 | 1.7 |
